# Highly efficient tamoxifen-inducible Cre recombination in embryonic, larval and adult zebrafish

**DOI:** 10.1101/2024.03.21.586128

**Authors:** Edita Bakūnaitė, Emilija Gečaitė, Justas Lazutka, Darius Balciunas

## Abstract

We have generated transgenic lines containing zebrafish-optimized CreER^T2^ recombinase under the control of a recombinant *ubb^R^* promoter consisting of the zebrafish ubiquitin promoter supplemented with an intronic enhancer from the carp *beta-actin2* gene. These lines enable highly efficient tamoxifen-inducible recombination in embryonic, larval and adult zebrafish.

**Abstract:** The ability to inactivate gene function in an adult organism is essential for studies of biological processes such as regeneration and behavior. This is best achieved by engineering an allele which could be conditionally inactivated using Cre recombinase and subsequently inactivating gene function using a drug-inducible Cre recombinase. Several recent studies clearly demonstrate feasibility of engineering such conditional alleles in zebrafish. Meanwhile, achieving sufficient degree of recombination to induce complete loss of function has remained a major limitation. Herein we address this limitation by engineering a recombinant ubiquitin promoter *ubb^R^* consisting of the zebrafish *ubiquitin* promoter supplemented with an intronic enhancer from the carp *beta-actin2* gene. Using phiC31-mediated targeted integration, we demonstrate that *ubb^R^* clearly outperforms both parental promoters as well as currently available ubiquitous CreER^T2^ driver lines at all embryonic and larval stages tested. Furthermore, the *ubb^R^:CreER^T2^*driver line we generated enables near-complete inactivation of floxed alleles in adult zebrafish hearts. Finally, we demonstrate that our *ubb^R^*promoter retains high activity when integrated at other genomic loci, making it uniquely suitable for robust expression of transgenes at all stages of zebrafish ontogenesis.

**Highlights:**

- Used targeted integration to directly compare different *CreER^T2^* drivers
- Generated a ubiquitous *ubb^R^:CreER^T2^* driver line capable of near-complete inactivation of floxed genes in adult zebrafish hearts
- Demonstrated that the recombinant *ubb^R^* promoter is suitable for robust transgene expression when integrated at different genomic loci

## Introduction

The Cre/lox recombinase system has been a staple of mouse genetics for more than a quarter century (Branda & Dymecki, 2004; Kühn et al., 1995; Rossant & Nagy, 1995; Zou et al., 1994). The system consists of two key components. The first component is a transgene expressing the recombinase in either native (Cre) or estrogen analog-inducible (CreER and derivatives) form under the control of either ubiquitous, inducible or tissue-specific promoter. The second component is a cassette containing at least two *loxP* sites. Cre-mediated recombination between the two *loxP* sites leads to either excision or inversion of the intervening DNA element depending on the nature and orientation of *loxP* sites.

A wide range of transgenic zebrafish lines expressing Cre and/or CreER derivatives under the control of ubiquitous and tissue-specific promoters have been generated by many different laboratories worldwide (reviewed in Carney & Mosimann, 2018). These “driver” lines are most commonly used in lineage tracing experiments where excision of the intervening DNA element results in permanent labelling of Cre-expressing cells and their progeny by fluorescent protein expression (Carney & Mosimann, 2018; Lalonde et al., 2022). While the mouse-optimized CreER^T2^ coding sequence is most widely used in zebrafish (Hans et al., 2011; Mosimann et al., 2011), use of mammalian codon-improved CreER^T2^ (iCre) (Tromp et al., 2023) and zebrafish codon-optimised CreER^T2^ (Kesavan et al., 2018b) resulted in improved recombination efficiency in zebrafish embryos.

The Cre/lox system has also been used to conditionally induce loss-of-function phenotypes in zebrafish. Both cassette excision and cassette inversion paradigms have been successfully deployed for this purpose (examples of these can be found in Burg et al., 2018; Grajevskaja et al., 2018; Gu et al., 2021; Hoshijima et al., 2016; Ni et al., 2012; Ogawa et al., 2021; Shin et al., 2023; Sugimoto et al., 2017; Trinh et al., 2011). To date, such studies have almost exclusively been carried at early stages of zebrafish development: only three studies have reported Cre-mediated conditional gene inactivation in adult zebrafish (Grajevskaja et al., 2018; Ogawa et al., 2021; Sugimoto et al., 2017).

All transgenes, including those regulated by apparently ubiquitous promoters, are subject to position effects, a phenomenon where the regulatory context of the genomic locus where the transgene is integrated exerts influence on transgene expression pattern and level (Hans et al., 2009, 2011; Lalonde et al., 2022; Mosimann et al., 2011). With the exception of enhancer trapping (Balciunas et al., 2004; Parinov et al., 2004; Scott et al., 2007), position effects are considered to be a detrimental side effect of transgenesis. Attempts to minimize position effects by incorporating border elements or insulators have had variable success (Caldovic et al., 1999; Grajevskaja et al., 2013). Probably the most reliable strategy to mitigate expression variability arising from position effects is to integrate transgenes into a predefined genomic locus (Lalonde et al., 2023; Mosimann et al., 2013; Roberts et al., 2014). This strategy has an additional advantage of generating a stable single-copy harbouring zebrafish line already at F_1_ generation, as opposed to Tol2-mediated transgenic zebrafish F_1_s that usually have multiple integrations of the transgene (Balciunas et al., 2006; Distel et al., 2009; Kawakami et al., 2004; Zhang et al., 2019).

The main goal for this study was to establish a ubiquitous 4-hydroxytamoxifen-inducible Cre driver line that enables efficient mutagenesis at all stages of zebrafish development as well as in adult organism. We compared the activity of three different promoter variants by integrating them into the same locus using phiC31 integrase and found that the recombinant *ubiquitin* promoter *ubb^R^* supplemented with a carp *beta-actin2* enhancer (Liu et al., 1990) significantly outperforms *ubb* and *actb2* promoters. Removal of bacterial vector backbone further improved activity, resulting in near-100% efficiency in larval zebrafish and adult hearts. Thus, our new ubiquitous CreER^T2^ driver line will be a valuable resource to investigators wishing to completely inactivate genes at various stages of ontogenesis. In addition, we demonstrate broad utility of our recombinant *ubb^R^* promoter by showing that *ubb^R^*-driven transgenes integrated into other genomic loci display similarly high activity.

Currently available ubiquitous CreER^T2^ drivers display diminished activity after 24 hours post-fertilization (Hans et al., 2011; Lalonde et al., 2022; Mosimann et al., 2011). To address this limitation, we engineered a recombinant *ubiquitin* promoter (*ubb^R^* henceforth) consisting of the zebrafish *ubiquitin* promoter (Mosimann et al., 2011) supplemented with an intronic enhancer from the common carp *beta-actin2* gene (Liu et al., 1990). The *ubb^R^* promoter as well as both parental promoters were cloned into a phiC31-mediated targeted integration vector to drive expression of zebrafish-optimized CreER^T2^* (Kesavan et al., 2018). phiC31-mediated targeted integration was selected instead of the widely used random integration using Tol2 transposon to minimize expression variability due to position effects and to quickly derive single-insertion lines. Bacterial vector backbone was flanked by FRT sites for excision using Flp recombinase. Transgenes were integrated into *tpl102Tg* docking site line using green-to-red eye color switch selection system (Roberts et al., 2014). We discovered that *ubb^R^:CreER^T2^\** performs exceptionally well across all stages of zebrafish development, and that removal of the bacterial vector backbone leads to additional increase in activity. Finally, we observe near-complete excision in adult hearts. To further test the utility of the *ubb^R^*promoter, we performed targeted integration into *tpl104Tg* docking site line and isolated a single-insertion random integrant using the *Sleeping beauty* transposon system (Davidson et al., 2003). Both transgenics displayed high activity across developmental stages, further confirming that *ubb^R^* promoter is suitable for robust expression of transgenes integrated into other loci.

## Results

### Variable performance of currently available ubiquitous CreER^T2^ drivers

While recombination of one allele in a subset of cells is sufficient for lineage tracing experiments, very high frequency of biallelic recombination is likely required to achieve loss-of-function phenotype for conditional mutants. Notably, using ubiquitous *ubb:CreER^T2^* driver line *Tg(−3.5ubb:CreERT2, myl7:EGFP)zf2148*, a single-copy sub line of *Tg(−3.5ubb:CreERT2, myl7:EGFP)cz1702* (Mosimann et al., 2011), induction of recombination by addition of 4-hydroxytamoxifen (4-HT) at 10 hours post fertilization - prior to known onset of *tbx20* expression - failed to induce a severe cardiac defect in zebrafish embryos homozygous for the floxed *tbx20^tpl145^*allele (Burg et al., 2018). We speculated that stronger CreER^T2^ drivers may yield more robust loss-of-function phenotypes and sought to compare the ability of *ubb*:*CreER^T2^* (Mosimann et al., 2011) to that of two heat-shock and 4-HT inducible *Tg(hsp70l:mCherry-T2A-CreER^T2^)* lines, designated #12 and #15 (Hans et al., 2011), (Figure 1A-B). Female fish heterozygous for the driver and homozygous for floxed *tbx20^tpl145^* allele were crossed to *tbx20^tpl145/tpl145^* males, and embryos were treated with heat-shock and/or 4-HT at indicated time points to induce excision of the exon 2 of the *tbx20* gene. Loss of *tbx20* function leads to significantly reduced cardiomyocyte numbers by impaired proliferation of heart muscle cells (Just et al., 2016) and leads to formation of a severe cardiac edema (Figure 1C). We observed major differences in the ability of different CreER^T2^ drivers to induce this loss-of-function phenotype.

**Fig 1.**
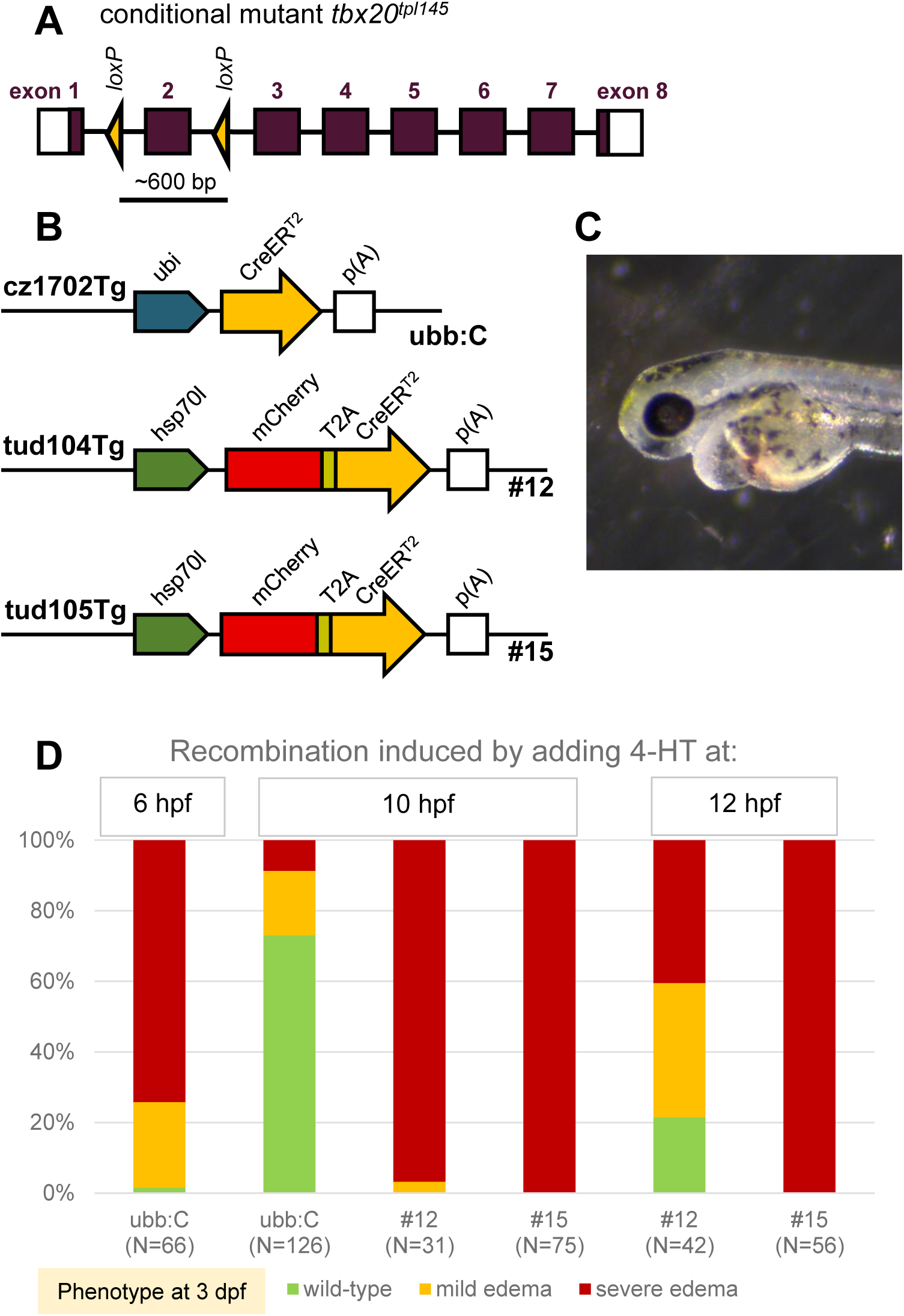
Non-uniform performance of currently available ubiquitous CreER^T2^ drivers. **A.** Diagram of the floxed tbx20 allele *tbx20^tpl145^*. **B.** Most widely used ubiquitous 4-HT and/or heat-shock-inducible Cre-driver lines. **C.** Loss-of-function phenotype observed after successful recombination in both alleles of the *tbx20* gene before the onset of the gene expression. **E.** Phenotypic analysis of 3-day-old embryos harbouring different ubiquitous Cre-drivers after the induction of recombination at 6 hpf, 10 hpf or 12 hpf.

Consistently with previous observations, effectiveness of the *ubb*:*CreER^T2^*driver diminished substantially as early as 10 hpf (Figure 1D). In contrast, both *hsp70l:CreER^T2^*drivers were able to induce complete loss-of-function phenotypes at 10 hpf and, for driver #15, even at 12 hpf. Notably, regardless of the driver used, *tbx20* mRNA was readily detectable at 24 hpf by *in situ* hybridization (data not shown).

Although heat shock inducible drivers ensure strong expression and recombination efficiency, there are several disadvantages related to their usage. First, administration of heat shock inevitably has a biological impact on an embryo (T. H. Ingalls et al., 1969; T. Ingalls & Murakami, 1962; Kimmel et al., 1988; Krone et al., 1997; Schirone & Gross, 1968). Second, activity of a driver depends on the exact conditions of the heat shock (Mottola et al., 2020), complicating reproducibility of experimental results from one laboratory to another. Third, these drivers do not induce well at late stages of development (Hans et al., 2011; Yoshinari et al., 2012). Therefore, there is a need for a strong ubiquitous tamoxifen-inducible Cre driver line that does not require the application of a heat-shock.

### Recombinant *ubiquitin* promoter *ubb^R^* outperforms other promoters

In our attempts to engineer a more reliable ubiquitous CreER^T2^ driver, we decided to use zebrafish codon-optimised CreER^T2^ (CreER^T2^*) (Kesavan et al., 2018b). We wanted to compare two widely used promoters: the zebrafish *ubiquitin* (*ubb*), (Mosimann et al., 2011), and the common carp *beta-actin2* (Clark et al., 2011; Gibbs & Schmale, 2000; Sivasubbu et al., 2006). The *ubb* promoter was modified to improve the splice acceptor site from AG/AT to AG/GT (see Materials and Methods). In addition, we engineered a recombinant zebrafish *ubiquitin* promoter by adding an enhancer element from the carp *beta-actin2* promoter (Liu et al., 1990) into the first intron (subsequently referred to as *ubb^R^* for *R*ecombinant, all constructs are show in Figure 2A). It is well known that the integration locus of the transgene can have an impact on its expression (Kalvaitytė et al., 2023; Kalvaitytė & Balciunas, 2022; Roberts et al., 2014). To avoid variability due to position effects (Hans et al., 2009, 2011; Kawakami et al., 2016; Mosimann et al., 2011; Roberts et al., 2014), we used a phiC31-mediated targeted-integration system (Roberts et al., 2014) (Figure 2B). Thus, we constructed three targeting vectors, containing *attB*-*mRFP* marker for phiC31-mediated transgenesis, the promoter being tested, and CreER^T2^*-coding sequence followed by the bovine growth hormone (bgh) poly(A) signal (Figure 2B). We also added *FRT* sites for subsequent removal of bacterial vector backbone that can have a silencing effect on the nearby transgene (Chen et al., 2004; Hlavaty et al., 2004; Lu et al., 2012; Lusky & Botchan, 1981; Valera et al., 1994).

**Fig 2.**
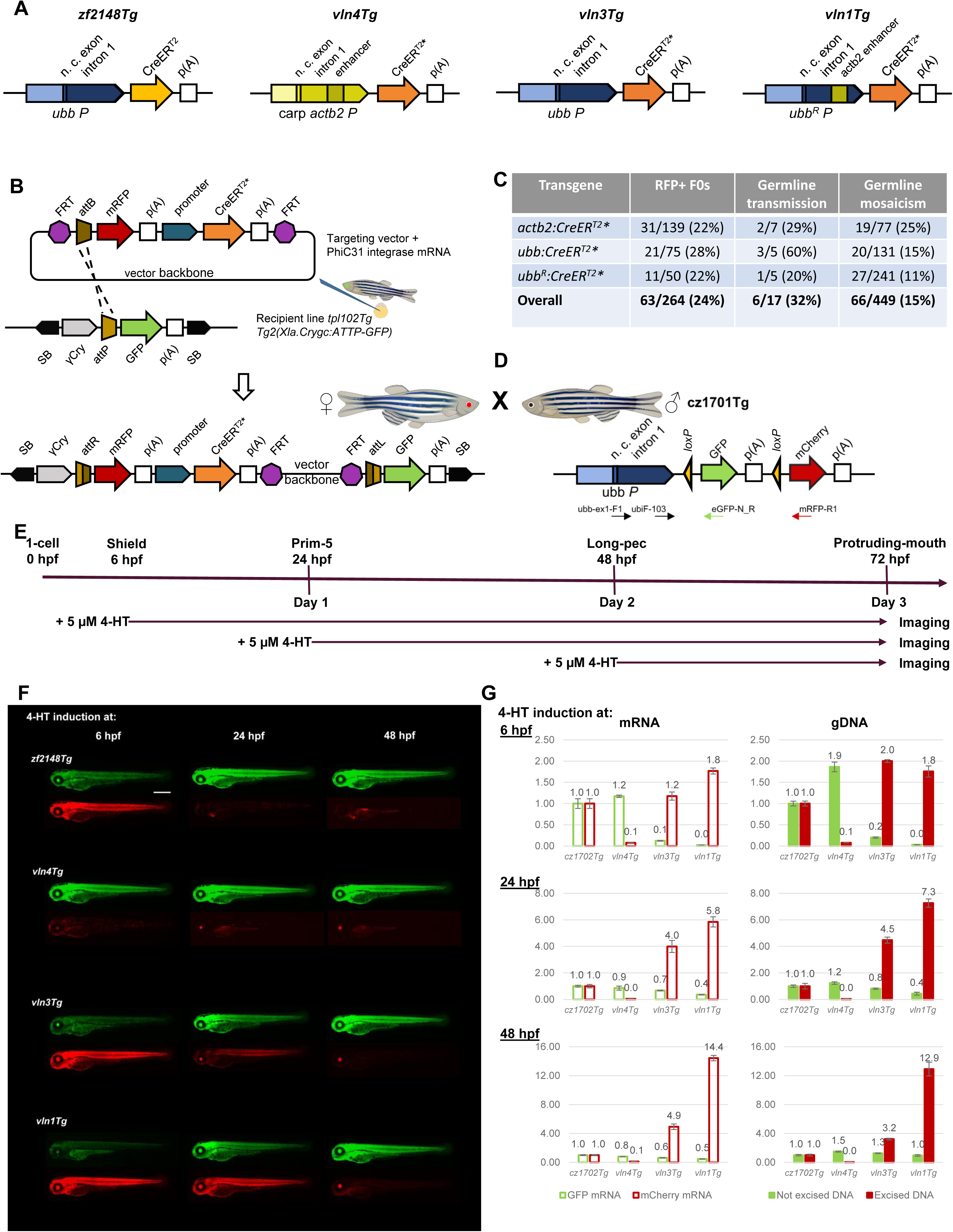
Construction and comparison of the efficiency of different ubiquitous 4-HT inducible Cre-drivers in zebrafish embryos and larvae. **A.** Diagrams of CreER^T2^ transgenes used in this experiment. *zf2148Tg* is a single-copy derivative of *cz1702Tg*, containing widely used mouse-optimised CreER^T2^ sequence under the control of zebrafish *ubiquitin* promoter. *vln4Tg* contains zebrafish-optimised CreER^T2^* sequence under the control of *beta-actin 2* promoter from common carp. *vln3Tg* contains zebrafish-optimised CreER^T2^* sequence under the control of zebrafish *ubiquitin* promoter. *vln1Tg* contains zebrafish-optimised CreER^T2^* sequence under the control of recombinant *ubb^R^* promoter, consisting of zebrafish *ubiquitin* promoter with carp *actb2* enhancer. **B.** Diagram of *phiC31*-mediated targeted integration system. *tpl102Tg* embryos containing an *attP* docking site have a green lens due to the *gamma-crystalline* promoter driving a GFP reporter. SB labels recognition sequences of Sleeping Beauty transposase used in generating the recipient transgenic line with docking site. Injection of a circular plasmid with *attB* and a red fluorescence reporter (targeting vector) into recipient line embryos leads to eye colour change upon *phiC31*-mediated integration in larvae. Reporter and CreER^T2^ transgenes are flanked with *FRT* recognition sequences for Flp recombinase, allowing subsequent removal of the vector backbone. **C.** Targeted transgenesis efficiency and germline mosaicism. **D.** Diagram of the ubiquitous lineage-tracing *ubi:switch* transgene used for the evaluation of the recombination efficiency of various transgenic Cre-driver lines. Horizontal arrows indicate the primers used for qPCR analysis. **E.** Experimental outline. Zebrafish males harbouring *ubi:switch* reporter were crossed to zebrafish females harbouring CreER^T2^ driver, embryos were treated with 5 µM 4-HT solution at 6, 24 or 48 hpf and kept in the dark until 3 dpf. At 3 dpf embryos positive for the CreER^T2^ driver were imaged and pools of 10 embryos were collected for quantitative analysis. **F.** Representative images of live 3 dpf embryos treated with 5 µM 4-HT solution starting at shield stage (6 hpf), prim-5 stage (24 hpf) or protruding mouth stage (48 hpf). After a successful recombination in the cell, the GFP-encoding DNA fragment is excised and *ubiquitin* promoter controls the expression of *mCherry*. Exposure of all red channel microscopy images was increased identically for easier evaluation. Scale bar, 500 µm. **G.** qPCR analysis of *GFP*/*mCherry* mRNA levels and unexcised/excised gDNA in 3-day-old larvae harbouring different CreER^T2^ transgenes after recombination induction at 6, 24 or 48 hpf. **B,D.** Zebrafish images were created in *biorender.com*.

For our experiments, we chose the recipient line *Tg(Xla.Crygc:attPGFP)tpl102* with *attP* docking site integrated into the chromosome 3. Fish with the docking site have a green lens due to the *gamma-crystallin* promoter driving a GFP reporter, and the integration of the targeting plasmid leads to eye colour change allowing selection of the transgenic F_0_ embryos. Upon injection of the targeting vectors mixed with phiC31 integrase mRNA into one-cell stage *tpl102Tg* zebrafish embryos, we observed that 22% (31/139), 28% (21/75), and 22% (11/50) of the embryos injected with *actb2:CreER^T2^\**, *ubb:CreER^T2^\**, and *ubb^R^:CreER^T2^\** encoding targeting vectors, respectively, were positive for lens RFP at 3 dpf. These embryos were raised to adulthood and outcrossed to wild-type fish to test if the integrated transgene is passed to F_1_ embryos. Germline transmission rates varied from 20% to 60% for different constructs. Altogether, 6 out of 17 outcrossed fish transmitted the transgene integrated into the docking site, with germline mosaicism varying between 11% and 25% (Figure 2C). Single-copy lines were designated *Tg*(*actb2:CreER^T2^*)vln4*, *Tg(ubb:CreER^T2^*)vln3*, and *Tg(ubb^R^:CreER^T2^*)vln1*.

Transgenic F_1_ females were then crossed to males homozygous for *Tg*(*–3.5ubi:loxP-GFP-loxP-mCherry*)*cz1701* (*ubi:switch*) reporter (Mosimann et al., 2011) (Figure 2D). Obtained embryos were divided into three separate groups and treated with 5 μM 4-HT at shield stage (6 hpf), prim-5 stage (24 hpf), and protruding-mouth stage (48 hpf). All the embryos were subsequently incubated in the dark until the analysis was performed at 3 dpf. At 3 dpf embryos positive for the CreER^T2^* driver (RFP signal in the lens) were selected, imaged under the fluorescent microscope to evaluate GFP and mCherry fluorescence, and pools of 10 embryos were collected for quantitative analysis of recombination efficiency (Figure 2E). For comparison, *ubb:CreER^T2^* (*zf2148Tg*, a single copy sub line of *cz1702Tg*) females were also crossed to *ubi:switch* males and the embryos were treated identically.

Our observations made with the *ubb:CreER^T2^* driver were entirely consistent with those reported previously (Lalonde et al., 2022; Mosimann et al., 2011). Recombination, as seen by induction of red fluorescence and concomitant reduction in green fluorescence, was strong after the 4-HT induction at 6 hpf, and diminished substantially when induced at later stages (Figure 2F). To our surprise, our transgenic line *vln4Tg* using the carp *beta-actin2* promoter to drive CreER^T2^* expression exhibited only negligible recombination of *ubi:switch* reporter at all time points tested. Our other two new transgenic lines, *vln3Tg* (*ubb:CreER^T2^\**) and *vln1Tg* (*ubb^R^:CreER^T2^\**), did not seem to offer a major improvement over *zf2148Tg* (*ubb:CreER^T2^*) when recombination was induced at 6 hpf, perhaps with the exception of some further reduction in green fluorescence. Both lines displayed a consistent increase in RFP fluorescence when induced at 24 hpf, and this increase was more pronounced for *vln1Tg*. When induced at 48 hpf (1 day before assessment), we did not observe any decrease in GFP fluorescence. RFP fluorescence was barely detectable for *vln3Tg* and more robustly detectable, but still not strong, for *vln1Tg*.

From each treatment group, three pools of 10 embryos were collected for quantitative assessment of recombination efficiency at the molecular level. We extracted total RNA and genomic DNA from these samples, used the RNA to synthesize cDNA, and performed quantitative PCR to assess relative recombination efficiency at the level of of mRNA (cDNA) and at the level genomic DNA (Figure 2G). qPCR results largely confirmed fluorescence-based observations. Relative to *zf2148Tg* (*ubb:CreER^T2^*), both *vln3Tg* and *vln1Tg* display an increase in excision of GFP, with ubb-GFP barely detectable at both mRNA (primers ubb-ex1-F1/eGFP-R) and gDNA (primers ubi-F103/eGFP-R) levels when *vln1Tg* driver was used. We also observed an up to 2-fold increase in ubb-mCherry for both drivers (primers ubb-ex1-F1/mRFP-R1 for mRNA and ubi-F103/mRFP-R1 for gDNA). Decreases in ubb-GFP are less pronounced at later stages, but increases in ubb-mCherry are very prominent, reaching greater than 10-fold improvement when induced at 48 hpf (Figure 2G).

### Removal of vector backbone increases recombination efficiency at late stages of development

We next tested if recombination efficiency can be further increased by deleting bacterial vector backbone sequences. This was achieved by the injection of Flp recombinase-coding mRNA into one cell stage embryos obtained from *vln1Tg* outcross (Figure 3A). Mosaic adults were then outcrossed to wild-type fish, resulting F_1_ embryos were grown to adulthood and genotyped for the removal of the vector backbone, and the resulting line was designated *Tg(ubb^R^:CreER^T2^*)vln2*. We then crossed *vln2Tg* males to *ubi:switch* and exposed the embryos to 4-HT at 6, 24 and 48 hpf to induce recombination.

**Fig 3.**
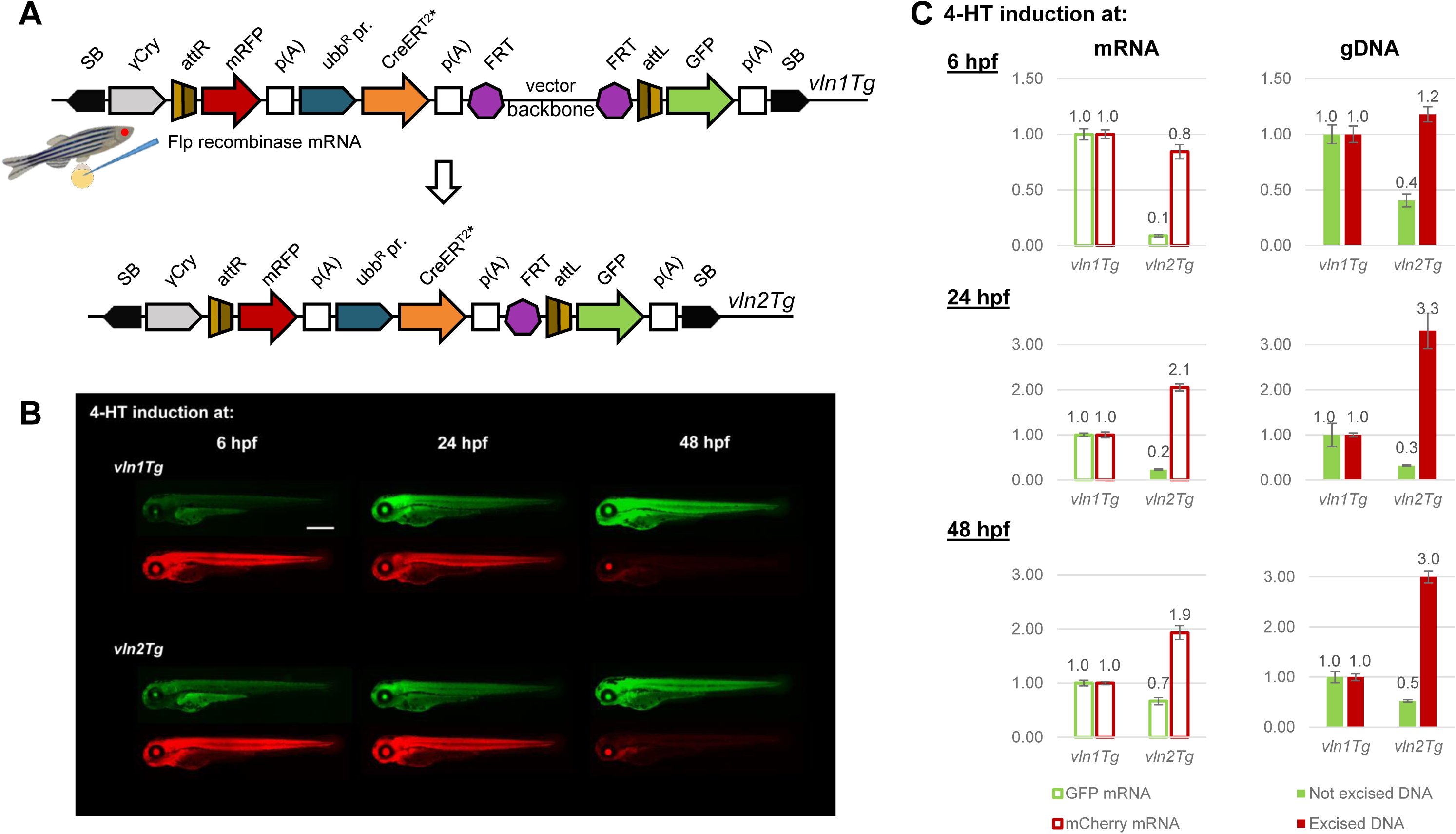
Vector backbone removal increases recombination efficiency at later stages of development. **A.** Diagram of vector backbone removal after injection of Flp recombinase-encoding mRNA into one-cell stage transgenic embryos. Zebrafish image was created in *biorender.com*. **B.** Representative images of live embryos, treated with 5 µM 4-HT solution at 6, 24 or 48 hpf, at 3 dpf. Exposure of all red channel microscope images was increased identically for easier evaluation. Scale bar, 500 µm. **C.** qPCR analysis of *GFP*/*mCherry* mRNA levels and unexiced/excised gDNA in 3-day-old larvae harbouring the transgene with vector backbone and after removal of the vector backbone.

GFP and mCherry fluorescence intensities of *vln2Tg* larvae treated with 4-HT at 6 hpf are similar to those of the parental strain *vln1Tg*, but *vln2Tg* larvae display more intense red fluorescence when treated at 24 and 48 hpf (Figure 3B). Quantitative PCR analysis also confirms that *vln2Tg* induces recombination more efficiently than *vln1Tg*: even though *vln1Tg* already reduced the GFP expression more than 100-fold, compared to *zf2148Tg*, at 6 hpf, removal of the vector backbone leads to further reduction at both mRNA and gDNA levels. We also observed 2-3-fold increase in ubb-mCherry signal at the mRNA and gDNA levels, respectively (Figure 3C). Therefore, we concluded that removal of the vector backbone has a positive impact on the expression of the transgene and, subsequently, the efficiency of recombination.

### *vln2Tg* zebrafish line enables efficient conditional mutagenesis during development of the zebrafish larvae

To evaluate the efficiency of our new *vln2Tg* driver in inducing loss-of-function phenotypes, we tested it on three different conditional mutants. Firstly, we wanted to see if this new driver can induce loss-of-function phenotypes of floxed *tbx20^tpl145^* zebrafish line (Figure 4A). We crossed *vln2Tg; tbx20^tpl145/tpl145^* females to *tbx20^tpl14/tpl1455^* males, then treated the resulting embryos with 5 µM 4-HT solution at 6 hpf, 10 hpf, and 24 hpf. At 3 dpf we collected the larvae with RFP signal in the lens (selection marker for *vln2Tg*), evaluated the development of the heart and collected pools of 10 larvae for quantitative analysis of recombination efficiency.

**Fig 4.**
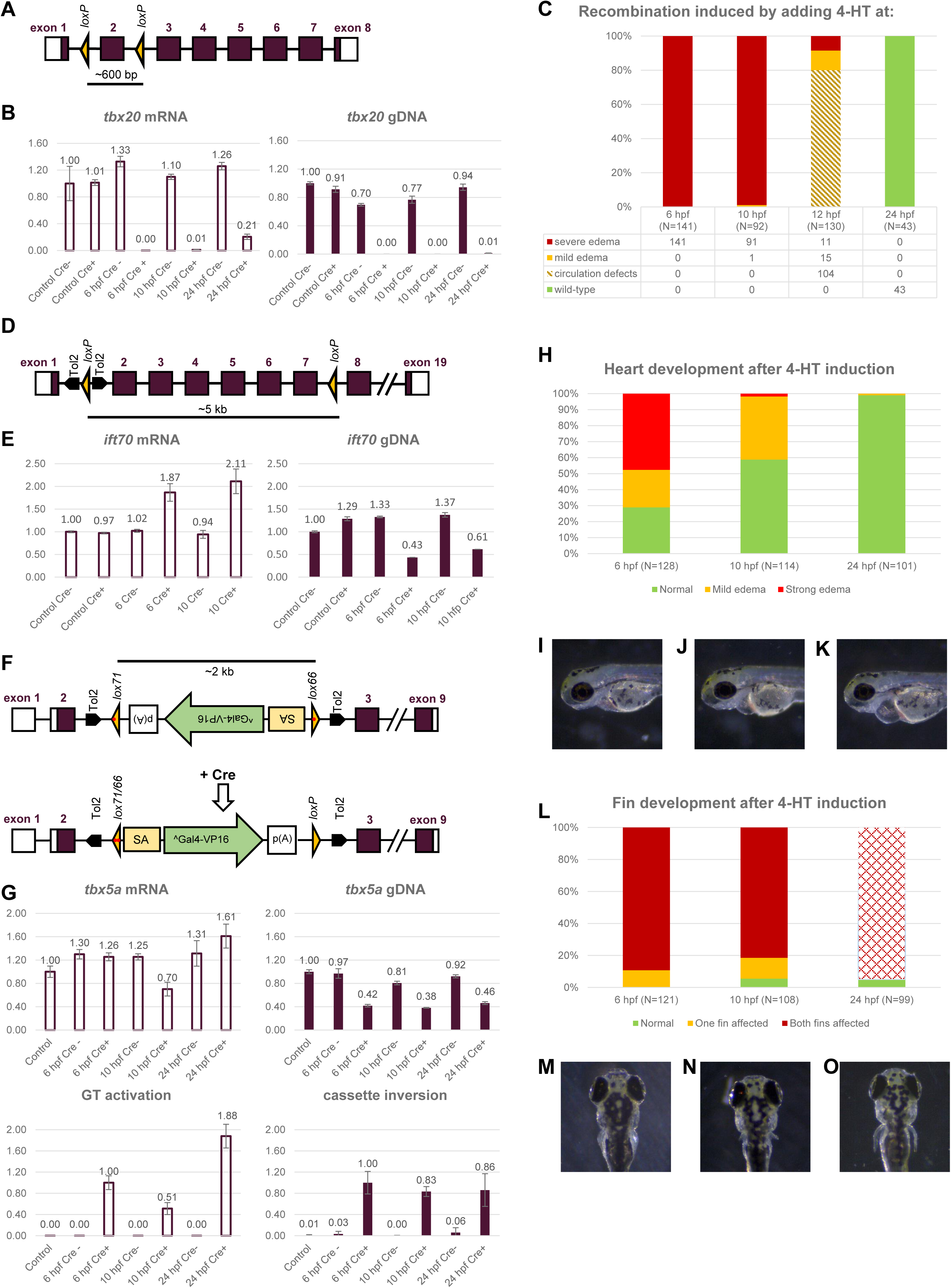
Recombination efficiency of three different conditional zebrafish mutants harbouring novel *ubb^R^:CreER^T2^* driver. **A.** Diagram of the floxed *tbx20* allele *tbx20^tpl145.^* **B.** qPCR analysis of the relative quantities of unexcised *tbx20* mRNA and *tbx20* gDNA levels after the induction of recombination at different time points during early development of the zebrafish embryos homozygous for *tbx20^tpl145^* allele. **C.** Phenotypic analysis of 3-day-old embryos homozygous for *tbx20^tpl145^*allele after the induction of recombination at 6 hpf, 10 hpf, 12 hpf or 24 hpf. **D**. Diagram of the floxed fleer allele *flr^tpl141^*. **E.** qPCR analysis of the relative quantities of unexcised *flr* mRNA and *flr* gDNA levels after the induction of recombination at different time points during early development of the zebrafish embryos homozygous for *flr^tpl141^*allele. **F.** Diagram of the *tbx5a* conditional gene trap allele. SA, carp beta actin splice acceptor; ^Gal-VP16, AUG-less Gal4-VP16; zp(A), zebrafish *beta actin* 3’ UTR and transcriptional termination sequences; cry, *X*. *laevis gamma crystallin* promoter; p(A), SV40 poly(A). The gene trap cassette is identical to that used in GBT-B1 gene trap vector (Balciuniene et al., 2013). **G.** qPCR analysis of the relative quantities of *tbx5a* mRNA and *tbx5a* gDNA levels and gene trap activation at mRNA level and cassette inversion at gDNA level after the induction of recombination at different time points during early development of the zebrafish embryos homozygous for *tbx5a^tpl58R^*allele. **H.** Heart development of the zebrafish embryos homozygous for *tbx5a^tpl58R^* allele after 4-HT induction at different developmental points. **I.** Wild-type zebrafish heart at 3 dpf. **J.** Mild heart edema of a 3-day-old zebrafish embryo observed after the inversion of the gene trap that turns off the expression of the *tbx5a* gene. **K.** Severe heart edema of a 3-day-old zebrafish embryo observed after the inversion of the gene trap that turns off the expression of the *tbx5a* gene. **L.** Pectoral fin development of the zebrafish embryos homozygous for *tbx5a^tpl58R^*allele after 4-HT induction at different developmental points. **M.** Pectoral fins of a 5-days-old zebrafish embryo with wild-type phenotype. **N.** 5-day-old zebrafish embryo with shortened pectoral fins observed after the inversion of the gene trap that turns off the expression of the *tbx5a* gene. **O.** Phenotype of a 5-day-old zebrafish embryo that was treated with 5 µM 4-HT solution at 24 hpf. The length of both fins are unaffected, albeit the shape of the fins differs from the embryo with wild-type phenotype.

qPCR analysis of gDNA revealed that excision of the floxed allele was almost complete at all time points tested. Compared to untreated larvae without a driver, relative quantity of the floxed allele dropped to 0%, 0%, and 1% after recombination was induced at 6 hpf, 10 hpf, and 24 hpf respectively (Figure 4B). RT-qPCR analysis of *tbx20* mRNA levels in the larvae indicate that after the 4-HT induction at 6 hpf and 10 hpf relative quantity of mRNA is equal to 0% and 1%, respectively. However, while qPCR analysis of gDNA shows that the floxed exon is excised almost completely when the embryos were treated with 5 µM 4-HT solution at 24 hpf, we still detected around 21% of wild-type mRNA in Cre-positive larvae. This mRNA should have been transcribed before the recombination was completed.

Almost complete excision of the exon 2 of the *tbx20* gene had a strong impact on cardiac development when recombination was induced at 6, 10 and 12 hpf. All (141/141) or nearly all (91/92) Cre-positive larvae displayed severe cardiac edema when induced at 6 hpf and 10 hpf, respectively. However, when the recombination was induced at 12 hpf, the observed phenotypes were different from those obtained using heat-shock and 4-HT inducible driver *Tg(hsp70l:mCherry-T2A-CreER^T2^)#15*. While after heat-shock treatment severe heart edema was observed in all (56/56) the embryos positive for the heat shock driver, *vln2Tg* induced severe or mild heart edema only in a subset of the embryos (11/130 and 15/130, respectively). Remaining embryos (104/130) had noticeable circulation defects and malformation of the heart. When the recombination was induced at 24 hpf, all the embryos (N=43) had a functioning heart and no edemas were observed.

We next tested our *vln2Tg* driver on floxed *ift70^tpl141^*zebrafish line (Burg et al., 2018). The main difference from *tbx20^tpl145^* is the distance between two *loxP* sites: ∼0.6 kb in *tbx20^tpl145^* vs ∼5 kb in *ift70^tpl141^* (Figure 4D). *Ift70* (intraflagellar transport 70, previously known as *flr*) gene encodes an essential regulator of cilia tubulin polyglutamylation, and mutant embryos exhibit ventral axis curvature, hydrocephalus, and kidney cysts (Pathak et al., 2007). Expression of the gene is first detected in Kupffer’s vesicle of seven somite embryos and the lateral mesoderm of 11 somite embryos (Pathak et al., 2007).

*vln2Tg* zebrafish females homozygous for *ift70^tpl141^*were crossed to *ift70^tpl141/tpl141^* males and the resulting embryos were treated with 5 µM 4-HT solution at 6 hpf or 10 hpf (before the onset of the *ift70* expression). At 3 dpf Cre-positive larvae were selected for phenotypical analysis and pools of 10 larvae were taken for quantitative evaluation of the floxed gene excision.

qPCR analysis showed that relative quantities of intact *ift70* gene sequence were reduced to 43% and 61% when recombination was induced at 6 hpf and 10 hpf, respectively, and phenotypical analysis of the larvae revealed that all the larvae had wild-type resembling phenotype. However, even though we observed a significant recombination efficiency, RT-qPCR analysis indicate that the expression of *itf70* is even higher in Cre-positive embryos. Compared to untreated embryos negative for Cre-driver, mRNA expression of *ift70* increases to 187% after the 4-HT treatment at 6 hpf, and to 211% after the 4-HT treatment at 24 hpf (Figure 4E).

Third zebrafish line used for the analysis of the efficiency of conditional mutagenesis induced by our Cre-driver is *tbx5a^tpl58t^* that has an invertible gene trap cassette flanked by LE/RE mutant *loxP* sites in the exon 2 of the *tbx5a* gene (Grajevskaja et al., 2018) (Figure 4F). Tbx5 is a highly conserved transcription factor known to be required for heart and upper limb development (Ahn et al., 2002; Garrity et al., 2002) with expression of the gene starting at gastrula stage (Tu et al., 2009).

*vln2Tg* zebrafish females homozygous for *tbx5a^tpl58R^* allele were crossed to *tbx5a^tpl58R/tpl58R^* males and the resulting embryos were treated with 5 µM 4-HT solution at 6 hpf, 10 hpf, and 24 hpf. Heart development was evaluated at 3 dpf, and development of pectoral fins was assessed at 5 dpf. Pools of 10 larvae sorted for RFP signal in the lens, indicating presence of the driver, were collected for quantitative analysis of gene trap inversion and *tbx5a* mRNA expression at 5 dpf.

Inversion of the gene trap results in exon 2 of *tbx5a* gene mRNA being spliced into to AUG-less Gal4-VP16 followed by zebrafish beta actin 3’ UTR and transcriptional termination sequences. This allows to quantify both the conditional mutant allele with inverted gene trap cassette and unrecombined *tbx5a* allele that ensures the expression of wild-type tbx5a. qPCR analysis revealed that after the induction of recombination at different developmental stages the relative quantity of wild-type *tbx5a*-expressing alleles are very similar and decreases to 42%, 38%, and 46%, when embryos are treated with 4-HT at 6, 10, and 24 hpf, respectively. To our surprise, RT-qPCR analysis did not detect a consistent corresponding decrease in wild type tbx5a mRNA levels (Figure 4G): after 4-HT induction at 6 hpf and 24 hpf relative levels of wt *tbx5a* mRNA increased to 125% and 161%, respectively. In contrast, *tbx5a* mRNA level decreased to 70%, however, when the recombination is induced at 10 hpf.

Nonetheless, phenotypic analysis of 4-HT treated zebrafish larvae performed at 3 dpf revealed that gene trap activation has a time-dependent impact on cardiac development (Figure 4H). After 4-HT treatment at 6 hpf, only 29% (37/128) of the driver-positive larvae had normally developed heart (Figure 4I), while 23% (30/128) and 48% (61/128) of larvae had mild (Figure 4J) or severe (Figure 4K) cardiac edemas, respectively. The percentage of larvae with normal heart increased to 59% (67/114) and 99% (100/101) after recombination induction at 10 hpf and 24 hpf, meanwhile only 2% (2/114) of larvae treated with 4-HT at 10 hpf had severe heart edemas, and none of the larvae treated at 24 hpf displayed severe heart defects (N=101) (Figure 4H).

Fin development was also impaired by the gene trap inversion (Figure 4L). When the recombination was induced at 6 hpf, none of the 5-day-old larvae displayed wild-type pectoral fins (Figure 4M): they all had defects of one (11%) or both (89%, Figure 4N) fins (N=121). Proportion of the impaired individuals was similar after the 4-HT treatment at 10 hpf: 6% (6/108) of larvae had normally developed fins, while 13% (14/108) and 81% (88/108) of larvae had one or both underdeveloped fins, respectively. It is interesting that while after 4-HT treatment at 24 hpf all larvae displayed pectoral fins of normal length at 5 dpf, 95% (94/99) of larvae had pectoral fins with posture different from wild-type resembling siblings (Figure 4O vs 4M). This suggests that *tbx5a* is important not only for the development, but also for the functioning of the pectoral fins.

### *ubb^R^:CreER^T2^\** driver is suitable for conditional mutagenesis in adult zebrafish

We next sought to investigate if *vln2Tg* can induce recombination in the hearts of adult zebrafish by testing this driver in the context of heterozygous *ubi:switch* reporter, homozygous floxed *tbx20^tpl145^*, homozygous floxed *ift70^tpl141^*, and homozygous conditional gene trap *tbx5a^tpl58R^*. We used adult females of 3 months of age or older and treated them with 5 µM of 4-HT solution three times for 24 hours in the dark, allowing them 24 hours of rest at normal husbandry conditions between each treatment. 7 days after the last 4-HT treatment the fish were euthanized, hearts were extracted and ventricles were collected for quantitative analyses (Figure 5A).

**Fig 5.**
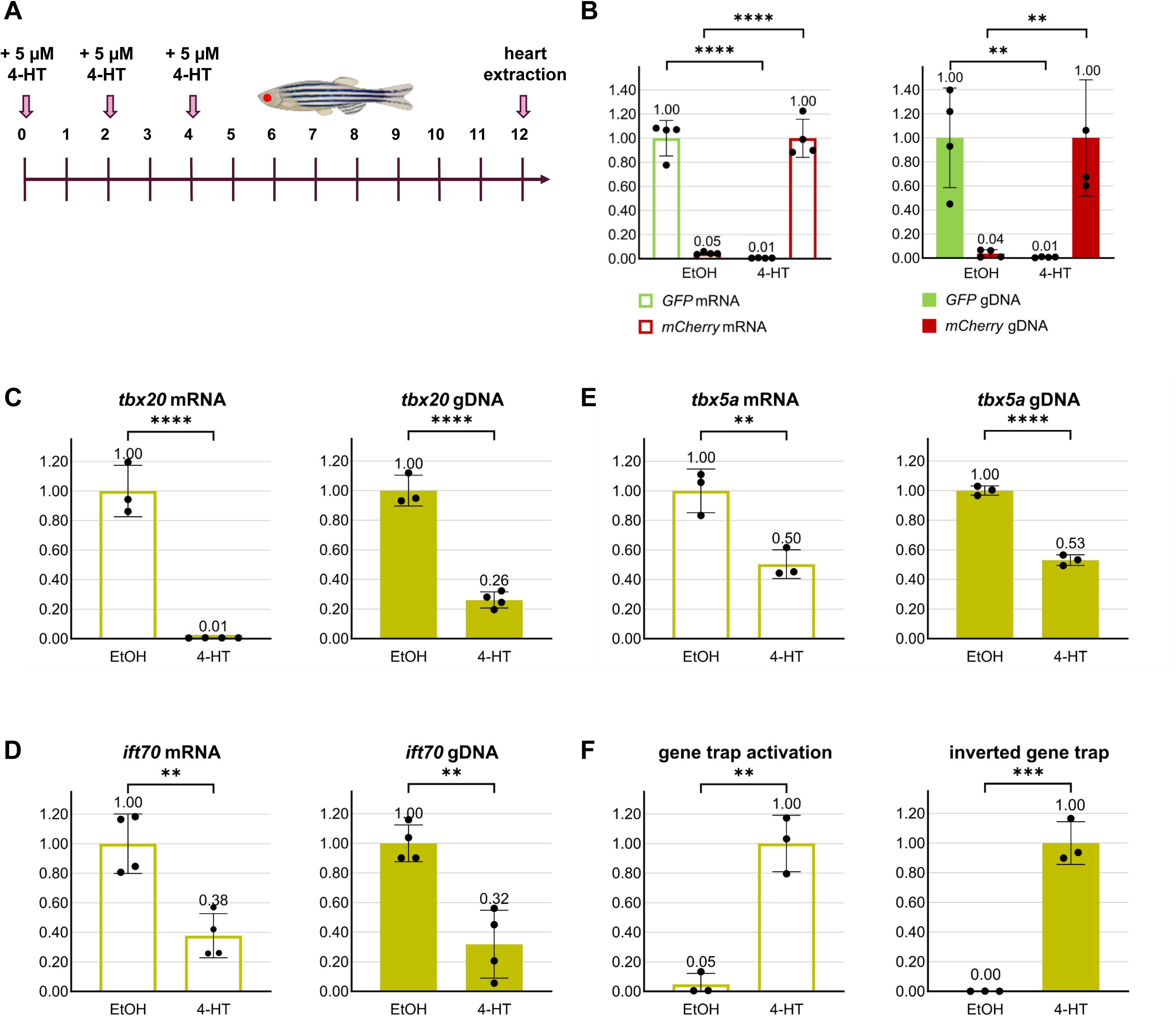
Recombination efficiency of adult zebrafish hearts harbouring novel *ubb^R^:CreER^T2^* driver. **A.** Experimental outline. Adult (3-month-old or older) zebrafish females were treated with 5 µM 4-HT solution three times for 24 h with one-day recovery period between treatments. Control fish were incubated in 0.01% ethanol (solvent for 4-HT) at the same conditions. One week after the last incubation, hearts were extracted and ventricles were used for further analysis. **B.** qPCR analysis of *GFP*/*mCherry* mRNA levels and unexiced/excised gDNA in heart ventricles of 4-HT or ethanol treated zebrafish with *ubi:switch* reporter. **C.** qPCR analysis of *tbx20* mRNA and gDNA levels in heart ventricles of 4-HT or ethanol treated *tbx20^tpl145^* zebrafish. **D.** qPCR analysis of *ift70* mRNA and gDNA levels in heart ventricles of 4-HT or ethanol treated *ift70^tpl141^* zebrafish. **E**. qPCR analysis of *tbx5a* mRNA/gDNA levels and gene trap activation/inversion in heart ventricles of 4-HT or ethanol treated *tbx5a^tpl58R^* zebrafish. **B-E**. n=3-4 biologically independent samples. Error bars, mean, s.d. Two-tailed Student’s *t*-test used to assess *P* values. *, *P*≤0.05; **, *P*≤0.01; ***, *P*≤0.001; ****, *P*≤0.0001.

The most efficient recombination is observed in floxed alleles with the *loxP* sites close to each other: according to RT-qPCR, after 4-HT induction, the relative quantity of ubb-GFP and exon 2 of *tbx20^tpl145^* mRNA drops to 1%, compared to that observed in ethanol-treated hearts (Figure 5C), even though approximately ¼ of the *tbx20^tpl145^* wild type allele remained refractive to excision at the gDNA level. Recombination efficiency of floxed *itf70^tpl141^* and gene trap of *tbx5a^tpl58R^* was noticeably lower. We were able to induce the excision of one third of the *itf70^tpl141^* allele and inversion of half of the *tbx5a^tpl58R^* allele, thus reducing the mRNA level of wild-type transcripts of these genes to 38% and 50%, respectively (Figure 5D-E).

We also noted a low level of Cre activity on the *tbx5a^tpl58R^* gene trap mutant and the *ubi:switch* reporter in the absence of induction by 4-HT. In both cases, we detect transcripts expected to arise after a successful recombination in ethanol-treated control hearts. The levels of these transcripts are 5% of what is observed in 4-HT-treated hearts (Figure 5B, F).

### *ubb^R^:CreER^T2^\** transgene displays high activity at other genomic loci

High activity of *Tg(ubbR:CreER^T2^*)vln1* could be attributed to inherent qualities of the transgene, serendipitously fortunate selection of the integration site (“safe harbor”, Kotin et al., 1992; Lalonde et al., 2022, 2023; Vooijs et al., 2001), or a combination of both factors. We therefore sought to test the activity of the *ubb^R^:CreER^T2^\** transgene at other loci by using a different docking site line and by performing a random integration using *Sleeping beauty* transposon (Balciunas et al., 2004; Davidson et al., 2003).

Using methodology described above for *tpl102*, we performed targeted integration of our cassette into *Tg(Xla.Crygc:attP-GFP)tpl104* (Roberts et al., 2014) and derived a sub-line in which the vector backbone was removed by Flp recombinase. The established zebrafish lines were named *Tg(ubb^R^:CreER^T2^*)vln7* and *Tg(ubb^R^:CreER^T2^*)vln8*, respectively. We have also subcloned the *ubb^R^:CreER^T2^\** into a *Sleeping beauty* (SB) transposon containing a lens-specific RFP transgenesis marker (Balciunas et al., 2004; Davidson et al., 2003) and generated a single-copy line *SB(ubb^R^:CreER^T2^*, Cry:RFP)vln10*. We then assessed the activity of these transgenes by crossing heterozygous transgenic female (*vln7Tg, vln8Tg*) or male (*vln10Tg*) fish to the *ubi:switch* reporter line, exposing embryos to 4-HT at 6, 24, and 48 hpf, and assessing fluorescence under a microscope at 3 dpf (Figure 6A). We noted that GFP and mCherry expression was quite similar when embryos were exposed to 4-HT at 6 hpf and 24 hpf, with *vln7Tg* appearing to display lowest levels of RFP fluorescence. In embryos exposed to 4-HT at 48 hpf, *vln10Tg* seemed to display highest level of RFP fluorescence (Figure 6A). We collected pools of 10 embryos exposed to 4-HT at 48 hpf and performed quantitative analysis of GFP excision and resulting mCherry expression at the mRNA and gDNA levels (Figure 6C-D). Quantitative analysis revealed that *vln7Tg* and *vln8Tg* display activity highly reminiscent of that of *vln1Tg* and *vln2Tg*, respectively, and that the activity of *vln10Tg* is similar to that of *vln2Tg*. Altogether, our data clearly demonstrate that newly-engineered *ubb^R^* promoter enables high level of CreER^T2^ activity when integrated into several different loci of the zebrafish genome.

**Fig 6.**
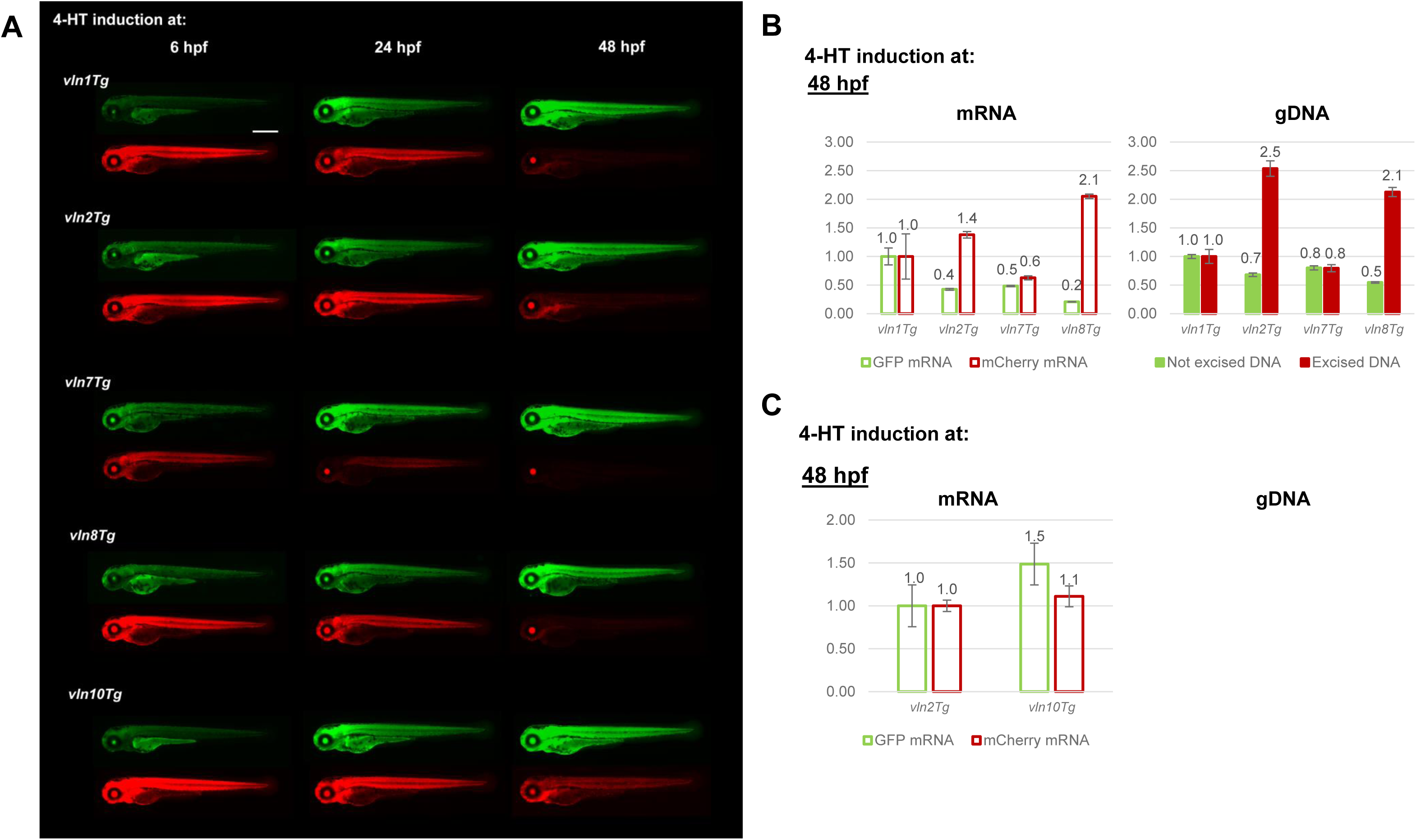
*ubb^R^:CreER^T2^\** transgene displays high activity at other genomic loci. **A.** Representative images of live 3 dpf embryos treated with 5 µM 4-HT solution at 6, 24 or 48 hpf. *vln1Tg*, *ubb^R^:CreER^T2^\** transgene with vector backbone at *tpl102Tg* docking site; *vln2Tg*, *ubb^R^:CreER^T2^\** transgene without vector backbone at *tpl102Tg* docking site; *vln7Tg*, *ubb^R^:CreER^T2^\** transgene with vector backbone at *tpl104Tg* docking site; *vln8Tg*, *ubb^R^:CreER^T2^\** transgene without vector backbone at *tpl104Tg* docking site; *vln10Tg*, *ubb^R^:CreER^T2^\** transgene integrated into a random genomic locus with SB100x transposase. Exposure of all red channel microscope images was increased identically for easier evaluation. Scale bar, 500 µm. **B.** qPCR analysis of *GFP*/*mCherry* mRNA levels and unexcised/excised gDNA in 3-day-old larvae with *ubb^R^:CreER^T2^\** transgene with and without vector backbone at two docking sites. **C.** qPCR analysis of *GFP*/*mCherry* mRNA levels and unexcised/excised gDNA in 3-day-old larvae with *ubb^R^:CreER^T2^\** at undetermined genomic locus after SB100x mediated random transgene integration.

## Discussion

This study describes generation and validation of a strong ubiquitous CreER^T2^* driver line that enables highly efficient 4-HT inducible recombination at all tested stages of zebrafish ontogenesis, thus providing a critical resource needed for studies of biological processes which occur late in development or in adults.

In addition to the two promoters widely used for quasi-ubiquitous transgene expression in zebrafish, zebrafish *ubiquitin* and common carp *beta-actin2*, we engineered a recombinant *ubb^R^* promoter consisting of the zebrafish *ubiquitin* promoter (Mosimann et al., 2011) and an intronic enhancer element from the carp *beta-actin2* (Liu et al., 1990). The activity of all three promoter variants was compared by performing phiC31-mediated targeted integration of otherwise identical transgenes driving zebrafish-optimized CreERT2* (Kesavan et al., 2018) into the *tpl102* docking site line (Roberts et al., 2014). Transgenic females were crossed to *ubi:switch* (Mosimann et al., 2011) males and recombination was induced by adding 4-HT at 6, 24 or 48 hours post fertilization. Recombination efficiency was assessed by fluorescence microscopy and quantitative PCR at 72 hpf.

To our surprise, activity of the carp *beta-actin2* promoter was barely detectable in our assay. Our findings contrast several previous observations noting strong expression in developing larvae as well as in adult zebrafish (Gibbs & Schmale, 2000; Sivasubbu et al., 2006). Some component of our transgenes must be rather specifically detrimental to the activity of the carp *beta-actin*2 promoter. We also noted relatively poor expression of the transgenesis marker, lens-specific RFP, in independently derived targeted integration lines containing the carp *beta-actin2* promoter, indicating that the whole transgene appears to be silenced. It is tempting to speculate that this transgene may be particularly sensitive to the presence of bacterial vector backbone.

Our observations made with the *ubiquitin* promoter are entirely consistent with the original study: recombination efficiency is very high when induced at 6 hpf, but diminishes significantly at 24 hpf and 48 hpf (Mosimann et al., 2011). Meanwhile, the recombinant *ubb^R^* promoter demonstrated improved recombination efficiency at all time points tested, including improvement by over an order of magnitude when induced at 48 hpf. Thus, although carp *beta-actin2* promoter itself does not seem to work well under our experimental conditions, it’s enhancer element is able to increase the activity of the zebrafish *ubiquitin* promoter.

PhiC31 integrates the whole plasmid, including the vector backbone, into the docking site locus. We flanked the bacterial backbone with FRT sites for removal using Flp recombinase. We were able to efficiently remove vector backbone, leading to further ∼2-fold increase in activity and leading to establishment of *vln2Tg* driver line used in subsequent experiments.

We next tested the ability of our *vln2Tg* CreER^T2^ driver line to induce *tbx20* loss-of-function phenotypes. Addition of 4-HT at 10 hpf leads to heart defects in 100% of the treated *tbx20^tpl145/tpl145^*embryos heterozygous for the driver, mimicking results obtained using heat-shock-inducible CreER^T2^ lines (Figure 1D). Induction of recombination at 12 hpf resulted in phenotypes less severe than those observed with heat-shock inducible driver: while the latter results in severe heart edema in all the treated embryos, the former mostly causes circulation defects and abnormal shape of the heart indicative of incomplete looping.

We next tested the ability of our driver to induce developmental phenotypes of a floxed *ift70^tpl141^* allele containing *loxP* sites spaced ∼5 kb apart. Excision of intervening DNA was not as efficient and did not exceed 60%. Consequently, we did not observe loss-of-function phenotypes. This observation taken together with recent publication failing to induce *tnnt2* loss-of-function phenotypes (Juan et al., 2024) suggests that minimizing distance between *loxP* sites may be a of critical importance when engineering floxed alleles.

Finally, we tested the ability of our driver to induce developmental *tbx5a^tpl58R^* LoF phenotypes using a conditional gene trap containing inverted *lox66*/*lox72* sites spaced ∼2 kb apart (Grajevskaja et al., 2018). We observed highly penetrant cardiac and pectoral phenotypes when induced at 6 hpf and milder phenotypes when induced at 10 hpf. Notably, Cre-mediated inversion of the gene trap did not significantly exceed 50%.

High activity of the *vln2Tg* driver at embryonic and larval stages prompted us to test if it would retain activity in adults. We tested it on all four target loci: *ubi:switch* (heterozygous), *tbx20^tpl145/tpl145^*, *tbx5a^tpl58/tpl58RR^*and *ift70^tpl141/tpl141^*. We observed near-complete recombination of *ubi:switch* and near-complete loss of *tbx20^tpl145/tpl145^*expression, indicating that *vln2Tg* driver is suitable for temporally-regulated inactivation of floxed alleles in adult hearts. Very high degree of excision of *tbx20^tpl145/tpl145^* at the level of genomic DNA - 74% - is particularly noteworthy. Only about 39% of all the cells in the adult zebrafish heart ventricle are cardiomyocytes (Patra et al., 2017), and not all cardiomyocytes express *tbx20*. Thus, our data clearly demonstrate that our driver is capable of inducing recombination at loci which are not transcriptionally active. In contrast, *tbx5a^tpl58/tpl58RR^* and *ift70^tpl141/tpl141^* were only “knocked out” at ∼50% and ∼30% efficiency, respectively (Figure 5D-E). We therefore propose that floxed genes with *loxP* sites in the near proximity to each other are superior to other types of conditional mutants. It is known that the distance between *loxP* sites is an important factor for the efficiency of recombination in other organisms (Araki et al., 1997; Coppoolse et al., 2005; Poulin et al., 2011; Zheng et al., 2000), and that modified *loxP* sites that are used in gene-trap conditional alleles are less effective than regular *loxP* sites (Araki et al., 1997; Yamauchi et al., 2022).

In this study, we used a docking site zebrafish line with pre-established genomic locus for transgene integration. This strategy allowed us to compare the efficiency of different promoters avoiding complications arising from position effects. The docking site line *tpl102Tg* (Roberts et al., 2014) used in our experiments proved to be a convenient tool for such experiments, considering high rate of RFP positive larvae after the injection of the targeting vector and PhiC31 mRNA, high germline transmission rate and germline mosaicism. Out data suggest that *tpl102Tg* line can be used as a safe harbour for efficient and reproducible transgenesis. To facilitate adoption of the promoter and the targeted transgenesis system, we constructed two targeted integration plasmids suitable for classical cloning and Gibson assembly: one for cloning any other transgene of interest under the *ubb^R^*promoter, and one for cloning any promoter of interest in front of the zebrafish-optimized *CreER^T2^\** (Figure 7).

**Fig 7.**
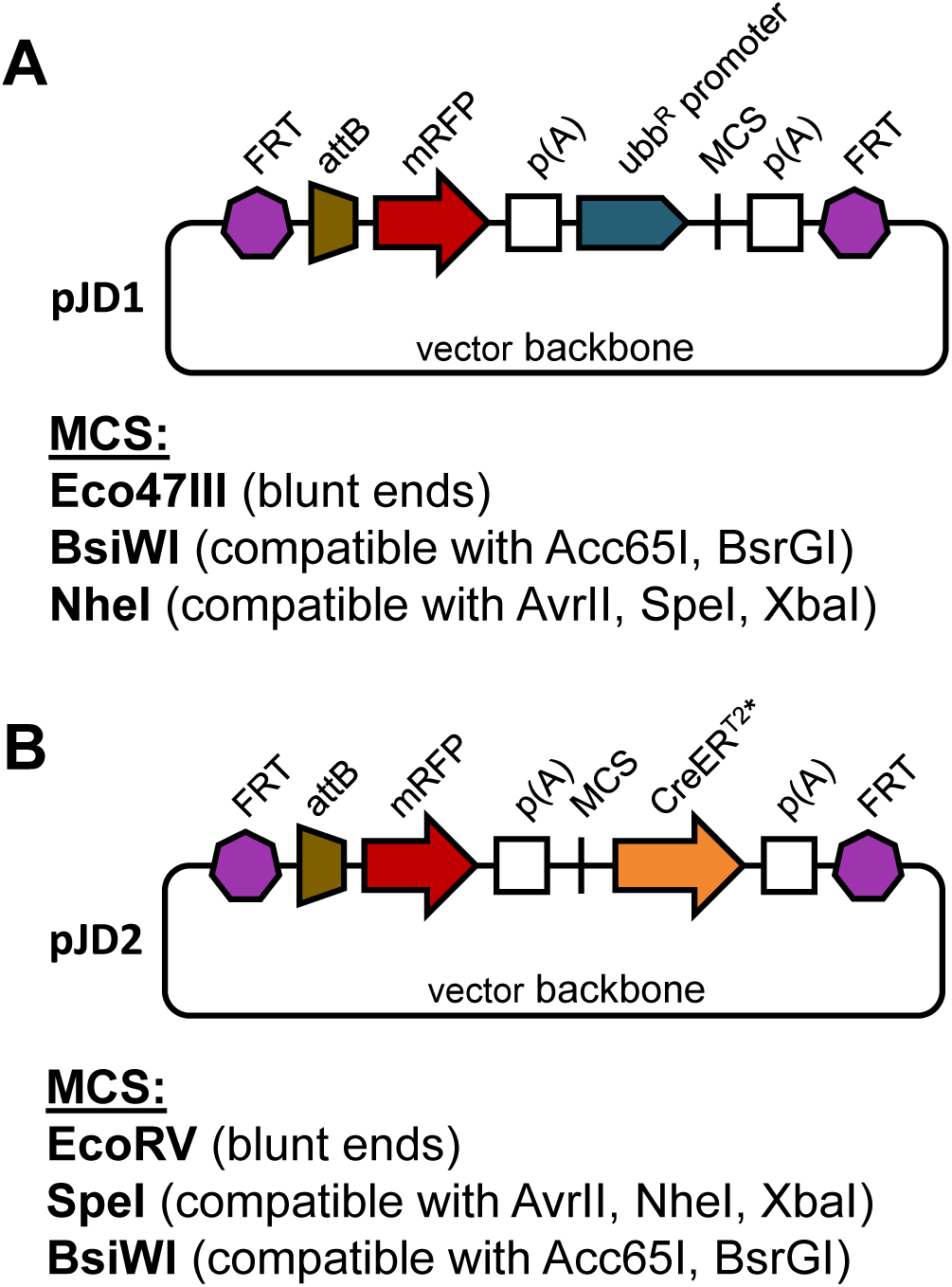
Vectors for targeted integration of transgenes. **A.** Plasmid pJD1, containing *attB*-mRFP marker for phiC31-mediated transgenesis, the *ubb^R^* promoter, and a multiple cloning site (MCS) for cloning a transgene of interest followed by the bovine growth hormone (bgh) poly(A) signal. **B.** Plasmid pJD2, containing *attB*-mRFP marker for phiC31-mediated transgenesis, a MCS for cloning the promoter of interest, and CreER^T2^*-coding sequence followed by the bovine growth hormone (bgh) poly(A) signal.

Finally, we demonstrate that *ubb^R^:CreER^T2^\** transgene is highly active when integrated into two additional genomic loci, indicating that the recombinant *ubb^R^* promoter will be tremendously useful for strong ubiquitous expression of other transgenes as well.

## Acknowledgements

This work was supported by European Social Fund (Project No. 09.3.3.-LMT-K-712-17-0014) under grant agreement with the Research Council of Lithuania (LMTLT). We are grateful to J. Dovidas for the construction of the plasmids pJD1 and pJD2; S. Hans and M. Brand for transgenic fish lines *tud104Tg*, *tud105Tg* and zebrafish-optimised *CreER^T2^*sequence; C. Mosimann for transgenic fish lines *cz1701Tg* and *cz1702Tg*; Z. Izsvák for SB100x transposase; P. Safabakhsh for construction of the pPS2 vector for SB100x transposase mRNA synthesis.

## Author Contributions

D.B. conceptualized and supervised the project. E.B. and E.G. constructed the plasmids. E.G. performed the microinjections, generated the transgenic fish lines. E.B. and E.G. collected embryo pools for quantitative analysis. J.L. performed imaging, image processing and statistical analysis. E.B. and J.L. performed quantitative analysis of recombination efficiency and evaluated loss-of-function phenotypes. E.B., J.L. and D.B. wrote the manuscript.

## Materials and Methods

### Zebrafish husbandry

Zebrafish (*Danio rerio*) stocks were kept under standard conditions at 27-27.5°C under 14:10 hour light:dark cycles, pH (7.4), and salinity-controlled conditions (Vet. Approval No. LT 59-13-002, LT 61-13-007). All experimental procedures were approved by the Lithuanian State Food and Veterinary Service (Approval No. G2-258). Male and female breeders from 3-12 months of age were used to obtain embryos and were used for all experiments with adult fish.

### Plasmid generation

All plasmids generated in this study were constructed using conventional restriction enzymes and CloneJet PCR Cloning Kit (Thermo Scientific™, K1231).

#### Plasmids used for targeted integration into *tpl102Tg* and *tpl104Tg* zebrafish lines

##### pEG10 actb2:CreER^T2^*

*attB* recombination site, *mRFP*, SV40 late poly(A) signal (Roberts et al., 2014); common carp *beta actin2* promoter (Gibbs & Schmale, 2000); zebrafish-optimised *CreERT2\** sequence (Kesavan et al., 2018a); bovine growth hormon *poly(A)* signal (Addgene #132775) cloned into *pUC18* vector (Addgene plasmid #50004) with *FRT* sites added at both ends of the whole construct.

##### pEG5 ubb:CreER^T2^*

*attB* recombination site, *mRFP*, SV40 late poly(A) signal (Roberts et al., 2014); zebrafish *ubiquitin* promoter (Mosimann et al., 2011) with improved splice acceptor site from AG/AT to AG/GT; zebrafish-optimised *CreERT2\** sequence (Kesavan et al., 2018a); bovine growth hormon *poly(A)* signal (Addgene #132775) cloned into *pUC18* vector (Addgene plasmid #50004) with *FRT* sites added at both ends of the whole construct.

##### pEG7 ubb^R^:CreER^T2^*

*attB* recombination site, *mRFP*, SV40 late poly(A) signal (Roberts et al., 2014); zebrafish *ubiquitin* promoter (Mosimann et al., 2011) with improved splice acceptor site from AG/AT to AG/GT and with enhancer element from common carp *beta actin2* promoter (Liu et al., 1990) inserted at *BsrGI* restriction site at forward orientation; zebrafish-optimised *CreERT2\** sequence (Kesavan et al., 2018a); bovine growth hormon *poly(A)* signal (Addgene #132775) cloned into *pUC18* vector (Addgene plasmid #50004) with *FRT* sites added at both ends of the whole construct.

#### Plasmid used for random integration of the transgene into wild-type fish

##### pEB19 ubb^R^:CreER^T2^*

zebrafish *ubiquitin* promoter (Mosimann et al., 2011) with enhancer element from common carp *beta actin2* promoter (Liu et al., 1990) inserted at *BsrGI* restriction site at forward orientation; zebrafish-optimised *CreERT2\** sequence (Kesavan et al., 2018a); SV40 late poly(A) signal, *X. laevis gamma-crystallin* promoter, and *mRFP* (Grajevskaja et al., 2018), cloned into *pT2 HB* (Addgene plasmid #26557).

#### Additional plasmids

##### pJD1 ubb^R^:GOI

*attB* recombination site, *mRFP*, SV40 late poly(A) signal (Roberts et al., 2014); zebrafish *ubiquitin* promoter (Mosimann et al., 2011) with improved splice acceptor site from AG/AT to AG/GT and with enhancer element from common carp *beta actin2* promoter (Liu et al., 1990) inserted at *BsrGI* restriction site at forward orientation; polylinker allowing integration of the gene of interest; bovine growth hormon *poly(A)* signal (Addgene #132775) cloned into *pUC18* vector (Addgene plasmid #50004) with *FRT* sites added at both ends of the whole construct.

##### pJD2 POI:CreER^T2^*

*attB* recombination site, *mRFP*, SV40 late poly(A) signal (Roberts et al., 2014); polylinker allowing integration of the promoter of interest; zebrafish-optimised *CreERT2\** sequence (Kesavan et al., 2018a); bovine growth hormone *poly(A)* signal (Addgene #132775) cloned into *pUC18* vector (Addgene plasmid #50004) with *FRT* sites added at both ends of the whole construct.

### Transgenic lines

#### The following published lines were used for this study

wild-type lines TL and AB obtained from the European Zebrafish Resource Center (EZRC) (https://www.ezrc.kit.edu/), *Tg(Xla.Crygc:ATTP-GFP)tpl102* (Roberts et al., 2014) and *Tg(Xla.Crygc:ATTP-GFP)tpl104* (Roberts et al., 2014)*, Tg(−3.5ubb:CreERT2, myl7:EGFP)zf2148* (Mosimann et al., 2011), *Tg(hsp70l:mCherry-T2A-CreER^T2^)tud104* (Hans et al., 2011)*, Tg(hsp70l:mCherry-T2A-CreER^T2^)tud105* (Hans et al., 2011)*, Tg*(*–3.5ubi:loxP-GFP-loxP-mCherry*)*cz1701* (Mosimann et al., 2011)*, tbx20^tpl145/tpl145^* (Burg et al., 2018), *ift70^tpl141^* (Burg et al., 2018), *tbx5a^tpl58RGt^* (Grajevskaja et al., 2018).

#### Zebrafish strains generated in this study

| Integration method | Construct | Abbreviation |
| --- | --- | --- |
| Targeted integration into *tpl102Tg* | *Tg(Xla.Crygc:ATTR-mRFP, actb2:CreER<sup>T2</sup>\*, Amp, ATTl-GFP)* | *vln4Tg* |
| Targeted integration into *tpl102Tg* | *Tg(Xla.Crygc:ATTR-mRFP, -3.5ubb:CreER<sup>T2</sup>\*, Amp, ATTl-GFP)* | *vln3Tg* |
| Targeted integration into *tpl102Tg* | *Tg(Xla.Crygc:ATTR-mRFP, ubb<sup>R</sup>:CreER<sup>T2</sup>\*, Amp, ATTl-GFP)* | *vln1Tg* |
| Targeted integration into *tpl102Tg* | *Tg(Xla.Crygc:ATTR-mRFP, ubb<sup>R</sup>:CreER<sup>T2</sup>\*, ATTl-GFP)* | *vln2Tg* |
| Targeted integration into *tpl104Tg* | *Tg(Xla.Crygc:ATTR-mRFP, ubb<sup>R</sup>:CreER<sup>T2</sup>\*, Amp, ATTl-GFP)* | *vln7Tg* |
| Targeted integration into *tpl104Tg* | *Tg(Xla.Crygc:ATTR-mRFP, ubb<sup>R</sup>:CreER<sup>T2</sup>\*, ATTL-GFP)* | *vln8Tg* |
| Sleeping Beauty-mediated random integration | *Tg(ubb<sup>R</sup>: CreER<sup>T2</sup>\*, Xla.Crygc:mRPP)* | *vln10Tg* |

### Transgenesis

Fertilised zebrafish eggs were generated by natural crossings. Targeted integration of the transgene into a docking site was achieved by injecting 3 nl of water solution containing 8.3 ng/μl plasmid DNA and 8 ng/μl of PhiC31-nos1-3ʹUTR integrase transcribed from pCS2+PhiC31onos1-3ʹUTR (as described in Roberts et al., 2014) into the yolks of one-cell stage *tpl102Tg* or *tpl104Tg* embryos. At 3 dpf embryos positive for RFP signal in the lens were selected and raised to adulthood. Mature fish were outcrossed to wild-type fish and their F_1_ progeny were screened for the transgenesis marker. Positive F_1_ fish were raised to adulthood and used for further experiments.

Vector backbone removal was achieved by injecting 3 nl of water solution containing 25 ng/μl of Flp° recombinase transcribed from pT3TS/Flp° (pDC50) (as described in Grajevskaja et al., 2018) into the yolks of one-cell stage embryos obtained from *Tg(ubb^R^:CreER^T2^*)vln1* and *Tg(ubb^R^:CreER^T2^*)vln7* outcross. F_0_ embryos were screened for RFP signal in the lens, raised to adulthood, outcrossed, and the resulting F_1_ fish were genotyped to confirm the excision of the vector backbone.

Random integration of the transgene was achieved by injecting 3 nl of water solution containing 8.3 ng/μl plasmid DNA and 30 ng/μl of Sleeping Beauty 100x integrase transcribed from pT3TS/SB100x (pPS2) (*in vitro* transcription was performed as described for pT3TS/Flp° (pDC50) in Grajevskaja et al., 2018) into the yolks of one-cell stage wild-type embryos. At 3 dpf embryos positive for RFP signal in the lens were selected and raised to adulthood. Mature fish were outcrossed to wild-type fish and their F_1_ progeny were screened for the transgenesis marker. Positive F_1_ fish were raised to adulthood, outcrossed to wild-type fish and their offspring were then used to establish single transgene insertion lines when transgene transmission of 50% or less was observed in the F_2_ generation. Mature F_2_ fish were used for further experiments.

### Heat shock and/or 4-HT treatment for CreER^T2^ induction in zebrafish embryos

4-hydroxytamoxifen (4-HT, H7904; Sigma) was dissolved in ethanol at a final stock concentration of 5 mM and kept in single-use aliquots in the dark at –80°C. To induce Cre activity in *ubiquitin*-based CreER^T2^ expressing embryos, 30-35 stage-matched embryos were transferred to a Petri dish with egg water that was freshly mixed with 4-HT heated for 10 min at 65°C to restore its activity (Felker et al., 2016) up to a final concentration of 5 µM. The treated embryos were put into a closed dark 28.5°C incubator and remained in 4-HT solution until imaging and sample collection at 3 dpf.

To induce Cre activity in *hsp70l*-driver CreER^T2^ expressing embryos, at 10 hpf pools of 50-60 embryos were transferred to a plastic tube with egg water preheated to 42°C with 5 μM 4-HT (for induction at 10 hpf) or without 4-HT (for induction at 12 hpf), kept in 42°C water bath for 3 minutes, then put into a closed dark 28.5°C incubator for 2 hours. After 2 hours embryos were transferred to a Petri dish (4-HT solution was added to the Petri dish with embryos to be induced at 12 hpf up to a final concentration of 5 μM), dead embryos were removed, and the remaining embryos were kept in the incubator until imaging and sample collection at 3 dpf.

### 4-HT treatment for CreER^T2^ induction in adult zebrafish

Mature (3-12 month-old) fish were incubated in 5 μM 4-HT solution three times for 24 h in the dark incubator with one-day recovery period between treatments (as described in Grajevskaja et al., 2018). One week after the last incubation, hearts were dissected, ventricles separated and frozen at –80°C for further analysis.

### Genotyping

Adult fish were genotyped by PCR using lysed tail clips. Primers used for genotyping are provided in table S1.

### Imaging and image processing

3-day-old zebrafish embryos were anesthetized in 0.04% (w/v) Tricaine (Sigma, E10521) solution. After anaesthesia they were mounted in a concave of a glass slide in 1% (w/v) low melting point agarose (Thermo Scientific™, R0801) ensuring their sagittal plane was parallel with the glass slide. Embryos were imaged using Leica DM5500B microscope with a HC PL FLUOTAR 5x/0.15 objective. Leica filter system TXR ET, k, Ex: 560/40 nm, Em: 630/75nm, dichroic: 585 nm was used to observe mCherry signal while filter system GFP ET, small, Ex: 470/40, Em: 525/50, dichroic: 495nm was used to observe eGFP signal. Fiji imaging software version 1.54h was used to pairwise stitch raw image files (Preibisch et al., 2009). The stitched images were cropped to a size of 2100 by 550 pixels and colored using “Merge channels” function of the same software. Raw image files are available upon request.

### Relative quantification of mRNA and gDNA levels

DNA and RNA were extracted from pools of 10 3 day-old embryos or ventricles of adult zebrafish hearts using TRI Reagent® (Sigma, T9424) followed by phenol-chloroform extraction. In brief, pools of embryos or heart ventricles were lysed and homogenized in 500 µl of TRI Reagent® using syringes with 21G or 23G needles, respectively. After addition of chloroform, shaking and centrifugation, phase separation was obtained. The top aqueous phase (containing RNA) was isolated and at least 400 ng RNA was used for subsequent cDNA synthesis (Maxima™ H Minus cDNA Synthesis Master Mix M1662, Thermo Scientific™) and RT-qPCR. The bottom organic phase (containing DNA) was washed with sodium citrate and subjected to ethanol purification to precipitate the DNA that was then dissolved in 8 mM NaOH and used for qPCR.

qPCR was performed in Rotor-Gene Q (QIAGEN) using Maxima SYBR Green qPCR Master Mix (Thermo Scientific™, K0253) without ROX passive dye. All reactions were performed in at least technical duplicates. Fold changes were calculated using the 2^-ΔΔCt^ method (Livak & Schmittgen, 2001). As a reference gene, eef1a1|1 was used for mRNA level quantification (McCurley & Callard, 2008) and aldh1a2 was used for gDNA level quantification (Burg et al., 2018). Primers used for qPCR are provided in table S2. All Ct values are listed in table S3.

### Statistics

Statistical analysis was performed using GraphPad Prism version 8.4. Unpaired Student’s t-test was used to determine statistical significance, p<0.05 was considered significant. Data are shown as mean ± S.D.

